# Genome-Wide Analysis of Facial Regionalization in Zebrafish

**DOI:** 10.1101/114801

**Authors:** Amjad Askary, Pengfei Xu, Lindsey Barske, Maxwell Bay, Paul Bump, Bartosz Balczerski, Michael A. Bonaguidi, J. Gage Crump

## Abstract

Patterning of the facial skeleton involves the precise deployment of thousands of genes in distinct regions of the pharyngeal arches. Despite the significance for craniofacial development, how genetic programs drive this regionalization remains incompletely understood. Here we use combinatorial labeling of zebrafish cranial neural crest-derived cells (CNCCs) to define global gene expression along the dorsoventral axis of the developing arches. Intersection of region-specific transcriptomes with expression changes in response to signaling perturbations demonstrates complex roles for Endothelin1 (Edn1) signaling in the intermediate joint-forming region yet a surprisingly minor role in ventral-most regions. Analysis of co-variance across multiple sequencing experiments further reveals clusters of coregulated genes, with in situ hybridization confirming the domain-specific expression of novel genes. We then performed mutational analysis of a number of these genes, which uncovered antagonistic functions of two new Edn1 targets, *follistatin a* (*fsta*) and *emx2*, in regulating cartilaginous joints in the hyoid arch. Our unbiased discovery and functional analysis of genes with regional expression in zebrafish arch CNCCs reveals complex regulation by Edn1 and points to novel candidates for craniofacial disorders.

**Summary Statement:** Using zebrafish to purify distinct groups of embryonic cells, Askary et al. have created a detailed map of how thousands of genes are deployed to shape the developing face.

## Introduction

The vertebrate facial skeleton is generated from CNCCs that populate a series of pharyngeal arches (Platt, 1893; Schilling and Kimmel, 1994). Signaling from endodermal and ectodermal epithelia, as well as from CNCCs themselves, establishes nested patterns of gene expression in arch CNCCs, in particular along the dorsoventral axis (Medeiros and Crump, 2012; Mork and Crump, 2015). CNCCs then progressively adopt a number of fates, including cartilage, bone, and ligament (Bronner and LeDouarin, 2012), with a subset of cells remaining as progenitors for later differentiation and possibly adult repair (Paul et al., 2016). The shapes and functions of distinct facial regions are inextricably tied to the selection of these cell fates and the subsequent growth and rearrangements of skeletal cells (Kimmel et al., 1998). The earliest fate adopted by arch CNCCs is cartilage, which occurs first in ventral-intermediate arch regions and then spreads to ventral and dorsal poles (Barske et al., 2016). Domain-specific differences in cartilage versus bone fates likely contribute to region-specific skeletal morphologies in the face. In dorsal and intermediate domains, early cartilage differentiation must be actively suppressed to ensure proper formation of joints and later-forming intramembranous bones (Askary et al., 2015; Nichols et al., 2016). Identifying the molecular differences that prefigure these regional cell fate choices and behaviors is key to unraveling how the facial skeleton is assembled.

Candidate-based approaches, as well as forward genetic screens in zebrafish (Piotrowski et al., 1996; Schilling et al., 1996), have identified key members of craniofacial signaling pathways and their downstream targets (Minoux and Rijli, 2010). Edn1 signaling is required for gene expression and subsequent skeletal patterning in the intermediate and ventral-intermediate regions of the arches, including the joint-forming domain (Kurihara et al., 1994; Clouthier et al., 1998; Miller et al., 2000; Ozeki et al., 2004; Sato et al., 2008; Gordon et al., 2013). The Bmp pathway has an overlapping function in patterning the lower face (Tucker et al., 1998; Bonilla-Claudio et al., 2012), although it appears to be preferentially required for gene expression in ventral-most arch regions (Alexander et al., 2011; Zuniga et al., 2011). In contrast, Jagged-Notch signaling is required to pattern dorsal arch CNCCs, at least in the hyoid and posterior mandibular arches of zebrafish (Zuniga et al., 2010; Barske et al., 2016). Downstream targets have also been identified, including Edn1 activation and Jagged-Notch inhibition of the Dlx3/4/5/6 family in ventral-intermediate CNCCs (Beverdam et al., 2002; Depew et al., 2002; Talbot et al., 2010; Zuniga et al., 2010), and Bmp regulation of Hand2 in ventral CNCCs (Thomas et al., 1998; Miller et al., 2003; Yanagisawa et al., 2003; Zuniga et al., 2011; Bonilla-Claudio et al., 2012). However, the extent to which Edn1 and Notch globally regulate dorsoventeral gene expression, and whether such regulation is always in the same direction, remains incompletely understood.

Recently, genome-wide expression profiling experiments in mice have identified stage-specific expression signatures of craniofacial compartments, such as the mandibular, maxillary and frontonasal prominences (Feng et al., 2009; Fujita et al., 2013; Brunskill et al., 2014). Similar studies have revealed genes regulated by Bmp4 (Bonilla-Claudio et al., 2012) and Dlx5/6 (Jeong et al., 2008). However, these studies relied on dissection of facial prominences rather than purification of the arch CNCCs that generate the facial skeleton. As the arches consist of not only CNCCs, but also endodermal and ectodermal epithelia and mesodermal cores, whether the identified genes were expressed in CNCCs was not always clear. In the current study, we use the nested expression of *hand2:GFP* and *dlx5a:GFP* transgenes along the dorsoventral axis to identify genes with domain-specific expression in CNCCs of the zebrafish mandibular and hyoid arches. By combining this domain-specific profiling with effects of altered signaling on arch CNCCs (Barske et al., 2016), we demonstrate global roles of Edn1 and Jagged-Notch signaling in establishing intermediate/ventral-intermediate and dorsal arch gene expression, respectively, yet only a minor role for Edn1 in the ventral-most arches. We then used gene editing to test the requirements for 12 previously uncharacterized domain-specific genes and found opposing requirements for *fsta* and *emx2* in coordinating skeletal development in the intermediate hyoid arch. Whereas *fsta,* a positive target of Edn1, was required to inhibit cartilage differentiation in the developing hyoid joint, *emx2,* a negative target of Edn1, was promoted cartilage differentiation at the connection points between individual hyoid cartilages. Thus, in addition to providing a global description of dorsoventral gene expression in arch CNCCs, these findings uncover a complex role for Edn1 in balancing skeletal differentiation in the intermediate arches.

## Results

### Generation of domain-specific arch transcriptomes by combinatorial transgene labeling

We have previously reported using dual labeling by *sox10*:DsRed and *fli1a*:GFP transgenes to purify all arch CNCCs from embryos at 20, 28, and 36 hours post-fertilization (hpf), followed by mRNA isolation, cDNA library construction, and deep sequencing (Barske et al., 2016). Here, we performed two additional replicates at 36 hpf to better define a minimum set of 472 arch CNCC-enriched genes. Next, we took advantage of the nested patterns of *hand2*:GFP and *dlx5a*:GFP transgenes to isolate distinct subsets of arch CNCCs along the dorsoventral axis at 36 hpf, followed by RNA sequencing (Figure 1A). Whereas fluorescence activated cell sorting (FACS) of *hand2*:GFP+; *sox10*:DsRed+ cells enriches for the ventral-most CNCCs of the arches, FACS of *dlx5a*:GFP+; *sox10*:DsRed+ cells enriches for a broader domain of ventral to intermediate arch CNCCs (and the otic vesicle). We then compared how RNA-sequencing data from different experiments align to the zebrafish genome. In our analysis, we included new *hand2*:GFP and *dlx5a*:GFP samples, as well as the previously described *sox10*:DsRed+; *fli1a*:GFP*+* cells from wild types, *edn1* mutants, *jag1b* mutants, and embryos with elevated Edn1 (*hsp70I:Gal4; UAS*:Edn1) and Notch (*hsp70I:Gal4*; *UAS*:Notch1a-ICD) signaling. For each experiment, the majority of reads (~80-90%) could be aligned to a unique position in the zebrafish genome, and the percentage of uniquely aligned reads was independent of the total number of reads for each particular sample (Figure 1B). The uniformly high percentage of uniquely aligned reads is a positive indication for the quality of the RNA sequencing data.

**Figure 1.**
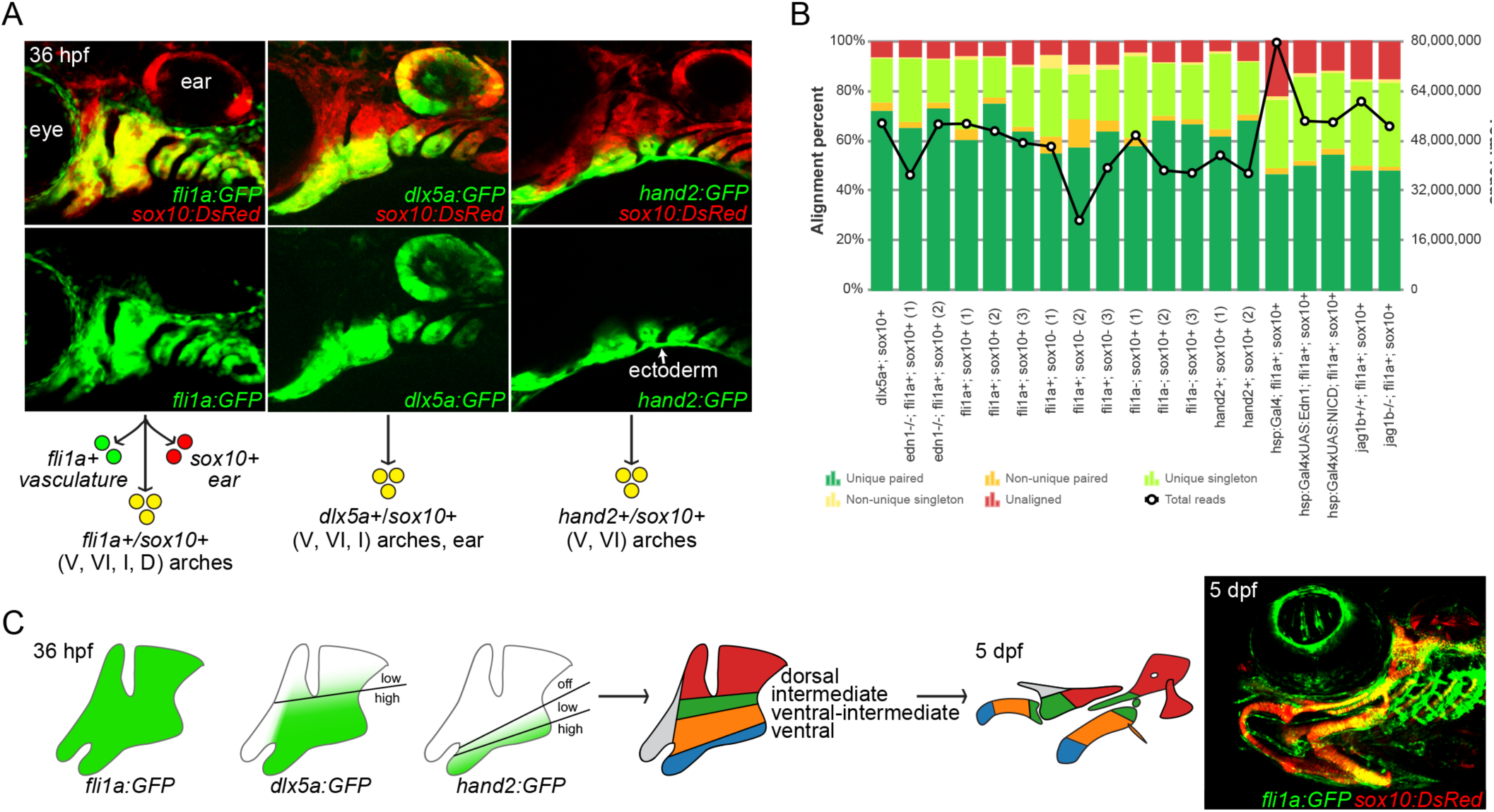
Isolation of arch CNCCs for RNA sequencing. (A) Cells were sorted using transgenic lines that label different populations of arch CNCCs. *fli1a*:GFP and *sox10*:DsRed overlap throughout arch CNCCs (yellow); *dlx5a*:GFP and *sox10*:DsRed in CNCCs of ventral (V) to intermediate (I) domains, as well as some dorsal (D) domain and the otic vesicle; and *hand2*:GFP and *sox10*:DsRed in ventral and more weakly ventral-intermediate (VI) CNCCs (modified from (Barske et al., 2016) and (Medeiros and Crump, 2012)). (B) Breakdown of alignment results for each RNA sequencing experiment, showing the percentage of reads aligned to a unique site in the genome and whether both paired end reads were aligned. The line graph shows the total number of reads acquired for each sample. (C) Dorsal, intermediate, ventral-intermediate, and ventral domains were defined by their relative levels of *hand2*:GFP and *dlx5a*:GFP. These domains map to specific parts of the larval craniofacial skeleton at 5 dpf (shown for context in a f/i1a:GFP; mx10:DsRed fish).

### Transcriptomic analysis reveals genes differentially enriched in *hand2*:GFP and *dlx5a*:GFP domains

In order to compare gene expression levels between samples, read counts were normalized to yield Transcript Per Million values (TPMs). We first identified the set of 472 genes enriched in arch CNCCs by filtering for genes with average TPMs greater than 2 across the three wild-type *fli1a*:GFP+; *sox10*:DsRed+ samples at 36 hpf, as well as enriched 1.5 fold or higher in double versus single positive cells (Supplementary Files 1 and 2). We then took advantage of the relative levels of *hand2*:GFP and *dlx5a*:GFP to subdivide arch CNCCs into four dorsoventral domains (Figure 1C). As previously described, *hand2*:GFP displays graded expression from strong in ventral-most domains to weaker in more ventral-intermediate regions, and *dlx5a*:GFP goes from strong in ventral to intermediate regions to weaker in dorsal domains (Medeiros and Crump, 2012). We therefore binned genes into clusters by comparing their relative expression in cells sorted with different transgenes: *hand2*:GFP^high^; *dlx5a*:GFP^high^ (“ventral”, 15 genes), *hand2*:GFP^low^; *dlx5a*:GFP^high^ (“ventral-intermediate”, 22 genes), *hand2*:GFP-; *dlx5a*:GFP^high^ (“intermediate”, 16 genes), and *hand2*:GFP-; *dlx5a*:GFP^low^ (“dorsal”, 30 genes) (see Table 1 for specific cut-off values). Confirming the validity of this filtering strategy, each dorsoventral cluster includes several genes with known expression in that domain (Supplementary Table 1). For example, the ventral cluster includes endogenous *hand2* (Miller et al., 2003) and homologs of genes known to be expressed in the ventral/distal domains of the murine arches (*foxf1* and *foxj2a*) (Jeong et al., 2004); the ventral-intermediate cluster includes endogenous *dlx5a,* as well as *dlx3b, dlx4b,* and *msxe* (Miller et al., 2000); the intermediate cluster includes *grem2b* (Zuniga et al., 2011) and genes required for joints such as *irx7* (Askary et al., 2015) and *nkx3.2* (Miller et al., 2003); and the dorsal cluster includes *jag1b* (Zuniga et al., 2010) and homologs of murine genes with dorsal arch expression (*pou3f3a* and *pou3f3b*) (Jeong et al., 2008).

**Table 1.**
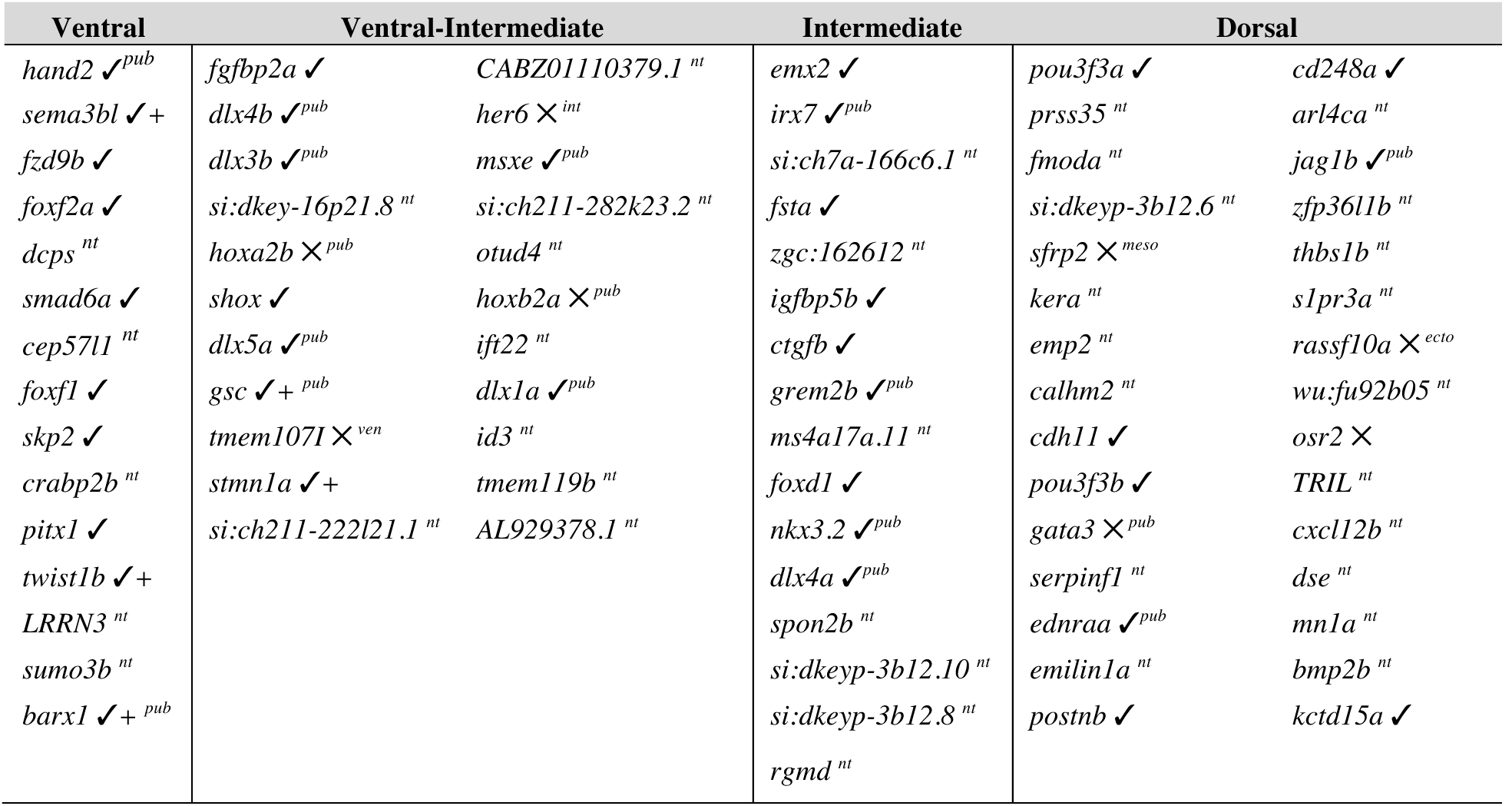
Predicted genes with dorsoventral-restricted arch expression. This table reflects expression in the mandibular and hyoid arches at 36 hpf only. **Ventral** genes were defined as showing TPM values in *hand2*:GFP+; *sox10*:DsRed+ cells >1.5 fold versus *fli1a*:GFP+; *sox10*:DsRed+ and >1.0 fold versus *dlx5a*:GFP+; *sox10*:DsRed+ cells. **Ventral-intermediate** genes were defined as TPM values in *dlx5a*:GFP+; *sox10*:DsRed+ cells >1.5 fold versus *fli1a*:GFP*+*; *sox10*:DsRed+ and between 1.0 and 3.0 fold greater versus hand2:GFP+; *sox10*:DsRed+ cells. **Intermediate** genes were defined as TPM values in *dlx5a*:GFP+; *sox10*:DsRed+ cells >0.67 fold versus *fli1a*:GFP+; *sox10*:DsRed+ and >3.0 fold versus *hand2*:GFP+; *sox10*:DsRed+ cells. **Dorsal** genes were defined as TPM values in *fli1a*:GFP+; *sox10*:DsRed+ cells >4.0 fold versus *hand2*:GFP+; *sox10*:DsRed+ and between 2.0 and 10.0 fold greater versus dlx5a:GFP+; sox10:DsRed+. ✓ positive expression in predicted domain; ✓ + positive expression in predicted domain as well as other domains; × expression outside the predicted domain; ^*ven*^ expression in the ventral domain; ^*int*^ expression in the intermediate domain; ^*meso*^ expression in mesoderm; ^*ecto*^ expression in ectoderm; ^*fn*^ expression in frontonasal NCCs; ^*pub*^ previously published; ^*nt*^ not tested.

### In situ validation of novel domain-specific gene expression in the arches

Given the inclusion of known genes with correctly predicted dorsoventral expression, we sought to validate the expression of uncharacterized genes in each domain-specific list. To do so, we conducted fluorescent in situ hybridization in 36 hpf embryos, along with *sox10*:GFPCAAX (membrane GFP) in a second color to highlight all arch CNCCs.

#### Ventral genes

Of the 8 predicted ventral genes tested, all showed some expression in the ventral mandibular or hyoid arches, yet their expression patterns were distinct (Figure 2A). Similar to *hand2* (Miller et al., 2000), we observed expression of *foxf1*, *foxf2a*, *fzd9b*, *smad6a*, and *skp2* in the ventral-most CNCCs of both arches, with *pitx1* showing more limited ventral expression in only the mandibular arch. In contrast, *sema3bl* and *twist1b* were expressed in the ventral domain and also to a lesser degree in subsets of dorsal arch CNCCs, reminiscent of the published expression pattern of *barx1* (Nichols et al., 2013; Barske et al., 2016), another gene on our ventral list. The weaker expression of these genes in dorsal relative to ventral arch CNCCs, together with their much lower levels in the intermediate domain, likely explains why they were enriched in the *hand2*:GFP data set and classified as ventral genes by our filtering scheme.

**Figure 2.**
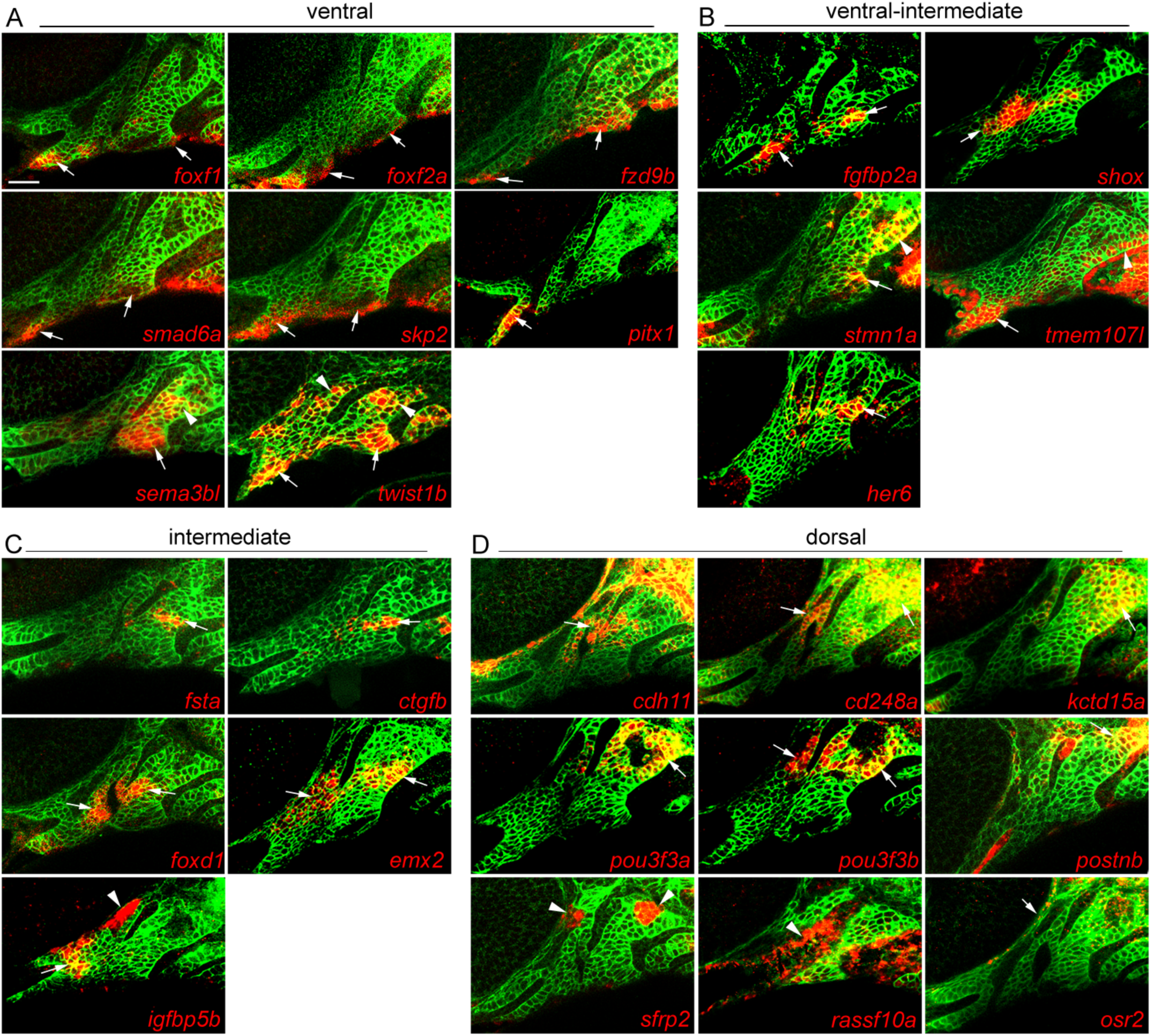
Arch expression of predicted domain-specific genes. Whole-mount fluorescent in situ hybridizations for select genes were performed in *soxl0*:GFPCAAX embryos at 36 hpf, with anti-GFP staining (green) showing CNCCs of the mandibular and hyoid arches. (A) *foxf1, foxf2a, fzd9b, smad6a* and *skp2* are expressed in ventral domains of both arches, *pitx1* only in the ventral mandibular arch, and *sema3b1* and *twist1b* in ventral and dorsal CNCCs. (B) *fgfbp2a, shox* and *stmn1a* are expressed in ventral-intermediate CNCCs, *stmn1a* in ventral-intermediate and dorsal CNCCs, *tmem1071* in ventral mandibular and posterior dorsal hyoid arches, and *her6* in a more dorsal domain. (C) *fsta, ctgfb, foxdl, emx2* and *igfbp5b* show specific intermediate domain expression; *igfbp5b* is also expressed in arch mesoderm. (D) *cdh11, cd248a, kctd15a, pou3f3a, pou3f3b,* and *postnb* are expressed in dorsal CNCCs, *sfrp2* in dorsal arch mesoderm, *rassf10a* in epithelia, and *osr2* between the dorsal first arch and the eye. Arrows point to expression in predicted domains, and arrowheads in other arch domains. Single optical sections are shown. Scale bar = 20 μm.

#### Ventral-intermediate genes

Among the predicted ventral-intermediate genes, two out of five tested showed expression within the ventral-intermediate domain (*fgfbp2a* and *shox*), one displayed both ventral-intermediate and some dorsal expression (*stmn1a*), one was expressed in the ventral mandibular arch and some dorsal posterior hyoid arch cells (*tmeml071*), and one in intermediate regions (*her6*) (Figure 2B). The ventral-intermediate list contains several known genes with apparent arch-wide expression (*hoxa2b* and *hoxb2a* (Hunter and Prince, 2002) and *dlx1a* (Sperber et al., 2008)) or ventral and dorsal expression domains (*gsc*) (Miller et al., 2000), although a closer examination of these previous reports suggests higher ventral-intermediate expression for some of these genes at 36 hpf.

#### Intermediate genes

Of the four previously characterized genes on this list, three are expressed in the intermediate joint-forming region and required for different joints of the face (*irx7*, *grem2b*, and *nkx3.2*) (Miller et al., 2003; Zuniga et al., 2011; Askary et al., 2015). The inclusion of *dlx4a* may reflect the broader expression of Dlx3-6 genes in both ventral-intermediate and intermediate domains (Talbot et al., 2010), although other members of this family were filtered into the ventral-intermediate category. All five newly tested genes showed highly specific expression in the intermediate domain, including *fsta, igfbp5b,* and *ctgfb* and homologs of mouse genes expressed in intermediate arch regions - *emx2* (Compagnucci et al., 2013) and *foxd1* (Jeong et al., 2004) (Figure 2C). Interestingly, *fsta* and *ctgfb* were largely restricted to the hyoid arch and *igfbp5b* to the mandibular arch.

#### Dorsal genes

We found 6 of 9 predicted dorsal genes to be enriched in the dorsal mandibular and hyoid arches: *cadherin11* (*cdh11*), *pou3fJ3a* and *pou3f3b* (homologs of mouse *Pou3f3* with dorsal expression (Jeong et al., 2008)), *cd248a, kctd15a* (also see (Gharbi et al., 2012), and *postnb* (Figure 2D). The other three genes were excluded from the *dlx5a*:GFP and *hand2*:GFP expression domains, as predicted, but they did not present typical “dorsal” expression patterns: *sfrp2* showed dorsal-specific expression that failed to co-localize with *dlx2a* (likely dorsal arch mesoderm), *rassf10a* was largely confined to the surface ectoderm rather than arch CNCCs, and *osr2* was expressed in only a few cells between the arches and the eye (Swartz et al., 2011). Although not tested here, previous reports also suggest dorsal-enriched expression of *ednraa* from 28-36 hpf (Nair et al., 2007; Zuniga et al., 2010) and expression of *gata3* in the more anterior maxillary prominence (Sheehan-Rooney et al., 2013b).

### Distinct regulation of domain-specific genes by Edn1 and Jagged-Notch signaling

Given their major roles in dorsoventral arch patterning, we next tested how Edn1 and Jagged-Notch signaling regulate the expression of genes in the dorsoventral domains defined by our RNAseq analysis. To do so, we intersected our previously published RNAseq data of gain or loss of Edn1 or Jagged-Notch signaling in arch CNCCs (Barske et al., 2016) with domain-specific genes identified based on enrichment in sorted *hand2*:GFP and *dlx5a*:GFP cells. We found that the expression of intermediate and ventral-intermediate genes, but not ventral and dorsal genes, was significantly reduced in *edn1*^-/-^ embryos (Figure 3A) and increased upon Edn1 misexpression (Figure 3B). In particular, intermediate genes were most affected by perturbation of Edn1 signaling, consistent with the sensitivity of intermediate skeletal elements (joints, symplectic, palatoquadrate) to partial reduction of Edn1 signaling in zebrafish (Walker et al., 2006). In contrast, loss of Jagged-Notch signaling in *jag1b*^-/-^ embryos resulted in a down-regulation of only dorsal genes (Figure 3C), and gain of Notch signaling resulted in a down-regulation of ventral, ventral-intermediate, and intermediate genes (Figure 3D). Gain of Notch signaling also had a trend towards increasing the expression of dorsal genes (Bonferroni corrected *p* = 0.18).

**Figure 3.**
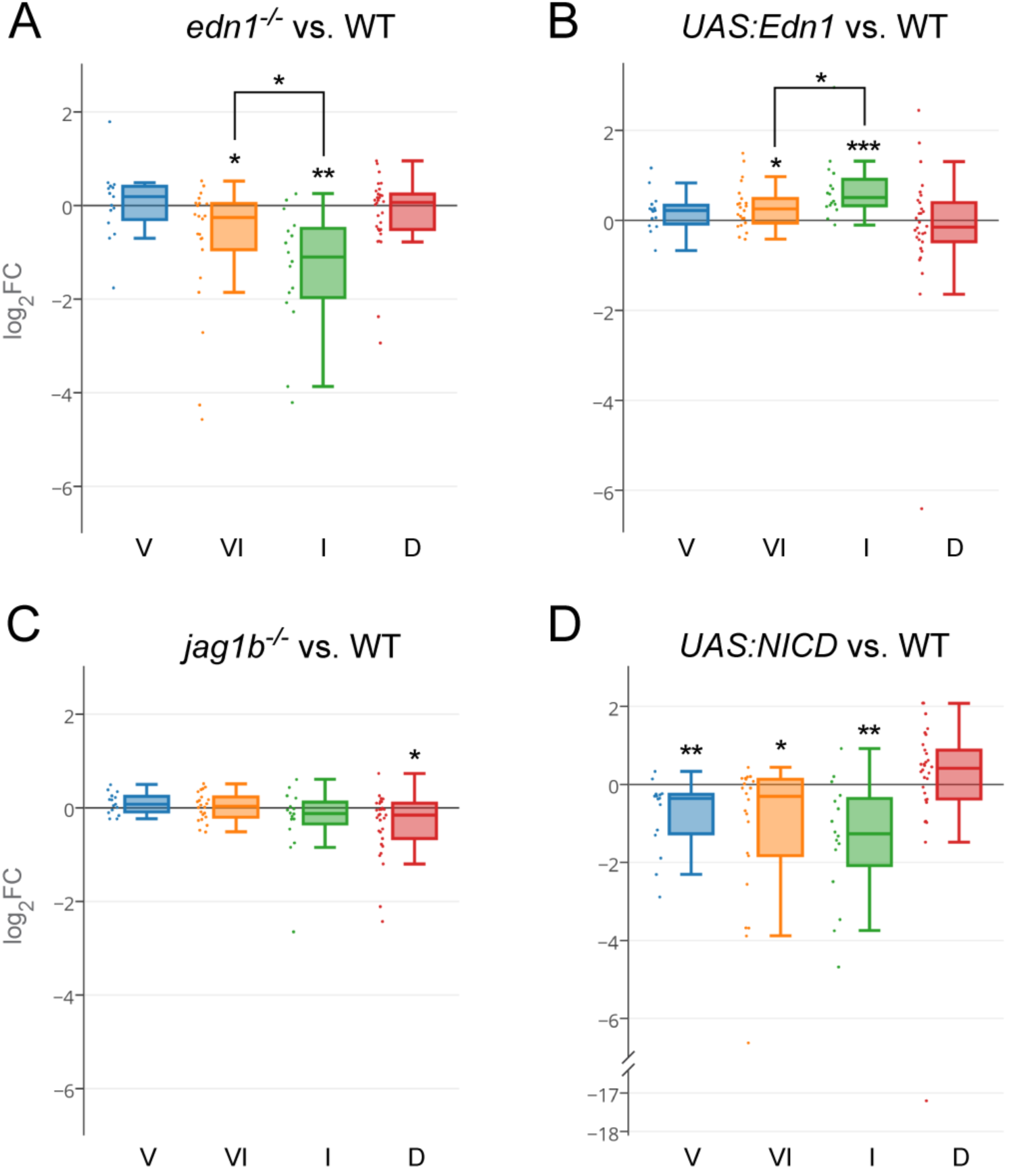
Domain regulation by Edn1 and Jagged-Notch signaling. (A) In *edn1*^-/-^ mutants, intermediate (I) domain genes are most downregulated, followed by ventral-intermediate (VI) genes. Ventral (V) and dorsal (D) genes are on average unaffected. (B) Edn1 overexpression results in greater upregulation of intermediate than ventral genes. (C) Dorsal genes are downregulated in *jaglb^-/-^* mutants. (D) Overexpression of the Notch intracellular domain (NICD) downregulates ventral, ventral-intermediate, and intermediate genes. See Materials and Methods for details of statistical analysis. * p < 0.05; ** p < 0.01; *** p < 0.001.

We next used in situ hybridization to confirm the predicted regulation of a subset of genes by Edn1 and Jagged-Notch signaling (Figure 4). Ventral-intermediate genes *fgfbp2a, shox,* and *stmn1a,* and intermediate genes *fsta* and *ctgfb,* were reduced in *edn1* mutants. However, intermediate gene *igfbp5b* was unaffected and intermediate gene *emx2* was variably upregulated in *edn1* mutants. Consistent with RNAseq data, ventral genes *smad6a, skp2,* and *fzd9b* were unaffected, and the ventral but not dorsal expression of *sema3b1* and the mandibular-specific ventral expression of *pitx1* were lost in *edn1* mutants. Of the dorsal genes examined, *pou3f3a* and *pou3f3b* were ventrally expanded in *edn1* mutants, *cdh11* expression shifted ventrally, and *cd248a* was largely unaffected. Reciprocally, *pou3f3a, pou3f3b,* and *cd248a* were reduced in *jag1b* mutants, with *cdh11* and *kctd15a* expression unaffected. In summary, we find intermediate and ventral-intermediate genes, but only a subset of ventral and dorsal genes, to be regulated by Edn1 signaling, and a subset of dorsal genes to be regulated by Jagged-Notch signaling.

**Figure 4.**
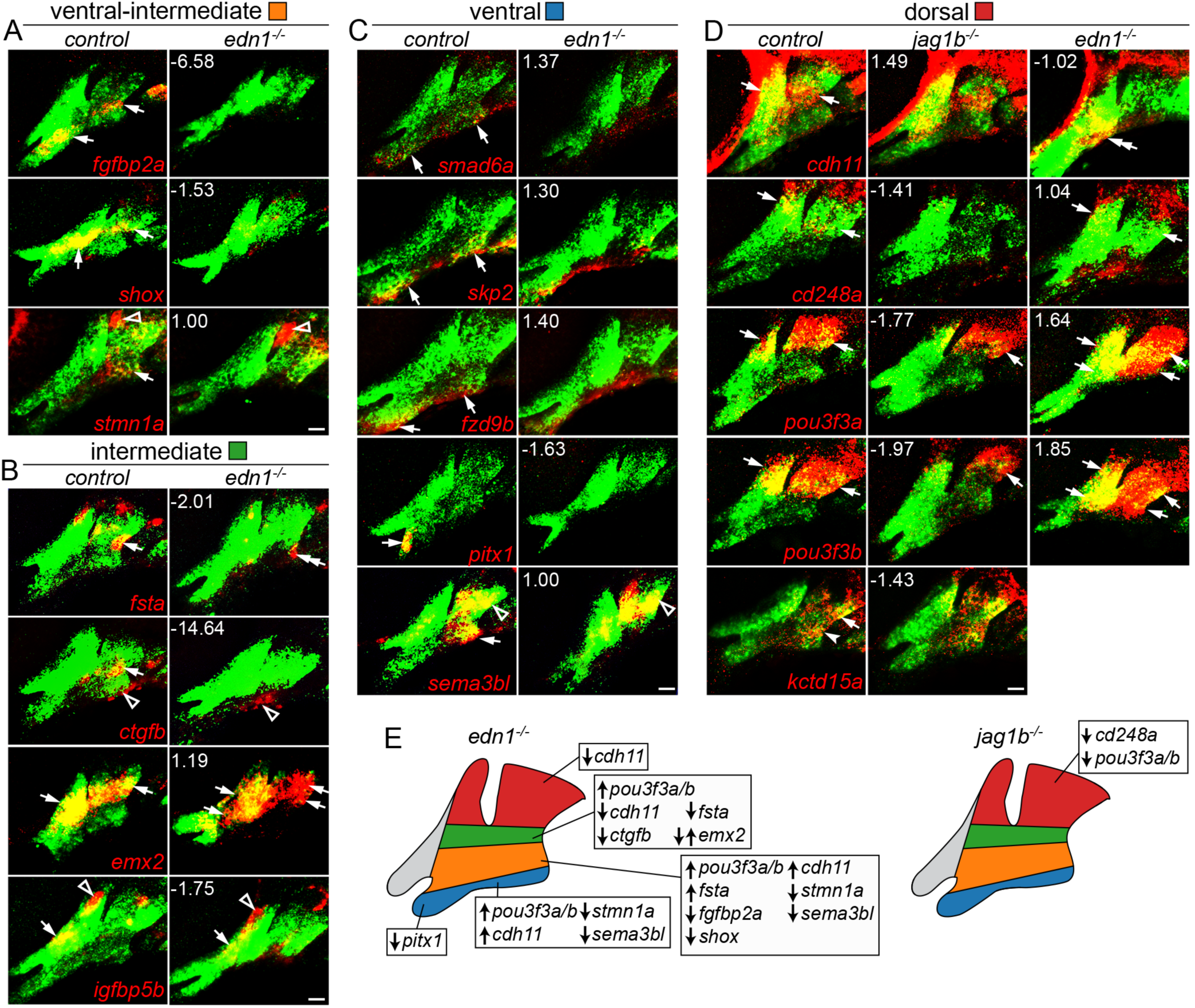
Changes in domain-specific gene expression in *edn1* and *jag1b* mutants. Two-color fluorescent in situ hybridizations were performed for genes of interest (red) and *dlx2a* (green) to label arch CNCCs at 36 hpf. Numbers indicate the gene expression fold-change in mutant versus wild-type *fli1a*:GFP+; *sox10*:dsRed+ CNCCs as determined by RNAseq. Maximum intensity projections show the mandibular (left) and hyoid (right) arches. (A) Ventral-intermediate expression of *fgfbp2a*, *shox* and *stmn1a* is lost in *edn1* mutants, yet pouch expression of *stmn1a* is unaffected. (B) In *edn1* mutants, intermediate expression of *fsta* and *ctgfb* is lost, *emx2* is variably upregulated (*n* = 3/10), and CNCC and mesoderm expression of *igfbp5b* is unaltered. We also note some ectopic ventral hyoid *fsta* expression in *ednl* mutants. (C) Ventral expression of *smad6a, skp2,* and *fzd9b* is normal in *edn1* mutants, yet ventral expression of *pitx1* and *sema3b1* is lost. Note that dorsal expression of *sema3bl* is unaffected. (D) In *jag1b* mutants, dorsal expression of *cd248a* is lost, *pou3f3a* and *pou3f3b* are reduced, and *cdh11* and *kctdl5a* are unaffected. In *edn1* mutants, *cdh11* expression shifts to ventral CNCCs, *pou3f3a* and *pou3J3b* are ectopically expressed in ventral CNCCs, and *cd248a* is largely unaffected. Arrows indicate expression in predicted arch domains, open triangles indicate additional expression domains, and double-arrows show expansion into other CNCC domains in mutants. Unless stated otherwise, consistent expression patterns were seen in a minimum of 3 wild types and 3 mutants for each experiment. Scale bar = 20 μm. (E) Summary of verified gene expression changes in *edn1* and *jag1b* mutants. Unaffected genes are not listed.

### Co-expression network analysis of pharyngeal arch genes

As an independent strategy to uncover genes co-expressed in arch domains, we performed a weighted gene co-expression network analysis (Zhang and Horvath, 2005) across 19 of our RNA-seq data sets (see Figure 1B). We limited this analysis to the 6000 genes exhibiting the greatest variance across all data sets and showing an expression level above 2 TPM in at least one experiment. A searchable dendrogram reflects the topological overlap metric (TOM), which is a measure of the correspondence in expression between genes across samples (Supplementary File 3). In order to determine the utility of TOM in uncovering novel genes within known networks, we examined five representative branches containing genes with validated dorsoventral-restricted expression (Figure 5A). Cluster 1 is composed of six genes, including the known dorsal gene *jag1b* (Zuniga et al., 2010). Of these, *cd248a*, *fgf20b* and *snail1a* were detected in dorsal arch CNCCs (Figure 2D and Figure 5B). Cluster 2 contains 14 genes, including a known intermediate gene (*grem2b*) required for joint formation in zebrafish (Zuniga et al., 2011), as well as four newly validated intermediate genes (*emx2, fsta, igfbp5b, foxd1*) (Figure 2C). This cluster also contains *twist1a,* which displays more complex expression in dorsal and ventral arch domains (Germanguz et al., 2007). Cluster 3 contains 11 genes, including five tightly clustered Dlx genes (*dlx3b, dlx4a, dlx4b, dlx5a, dlx6a*) known to be co-expressed in the ventral-intermediate domain (Talbot et al., 2010), as well as another known ventral-intermediate gene (*msxe*) (Miller et al., 2000) and a non-coding RNA (*si:ch673-351f10.4*) in an analogous position to the mouse *Evf2* gene, an antisense transcript that promotes the expression of the *Dlx5-6* locus (Feng et al., 2006). This cluster also contains an uncharacterized gene, *fgfbp2b,* which we find to be expressed in a subset of ventral-intermediate first arch CNCCs (Figure 5B). We also examined two distinct branches containing ventral-restricted genes. We verified five out of six genes in Cluster 4 as being restricted to the ventral-most arches (*pitx1, fzd9b, foxf1, foxf2a*) or expressed more strongly in the ventral arches (*sema3bl*) (Figure 2A). Cluster 5 contains a known ventral-restricted gene (*satb2*) (Sheehan-Rooney et al., 2013a) that tightly co-varies with *mitochondrial ribosome recycling factor* (*mrrf*), which we find to have similar ventral-restricted expression (Figure 5B).

**Figure 5.**
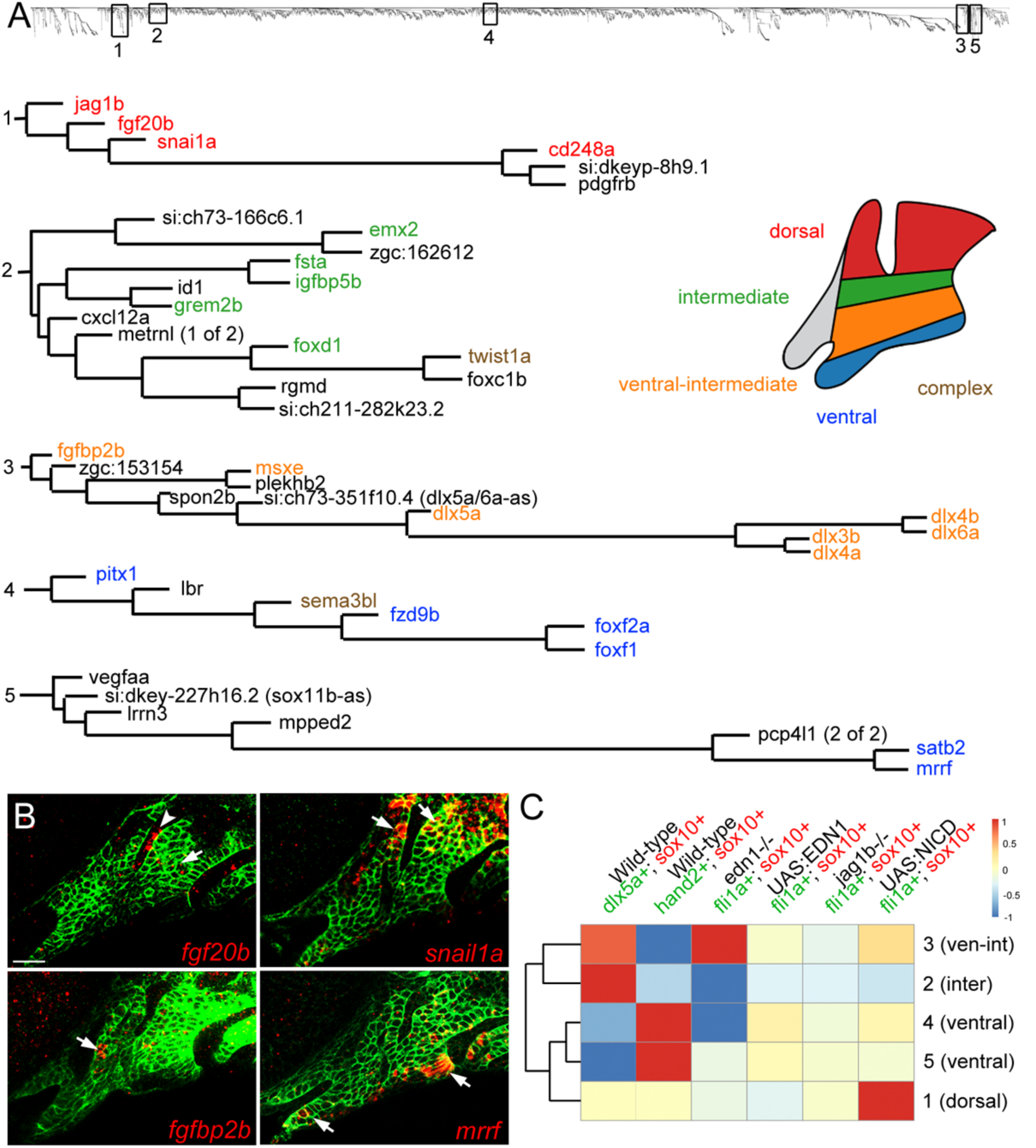
Co-variance network analysis reveals cohorts of similarly regulated arch genes. (A) Five representative clusters (1-5) were chosen from the dendrogram (top) generated by co-variance analysis. Gene names are color-coded based on expression patterns that are published or verified in this study. “Complex” refers to genes with broader expression in multiple domains. (B) Four genes discovered by co-variance analysis were confirmed by in situ hybridization (red) of soxiO.GFPCAAX embryos at 36 hpf; anti-GFP staining (green) marks CNCCs of the mandibular and hyoid arches. Arrowhead indicates *fgf20b* expression in the first pharyngeal pouch. Scale bar = 20 μm. (C) TDA analysis shows experiments that drove clustering (red) or disrupted clustering (blue). The dendrogram on the left shows the relatedness of clusters based on which data sets drove their clustering.

We next asked which RNAseq experiments drove gene co-regulation by iteratively computing the average TOM disruption caused by removing experimental groups, thus producing a TOM driver score for each experiment (Figure 5C). Expression in *dlx5a*:GFP+ cells was the strongest driver for Cluster 2, containing known and validated intermediate genes, and a strong driver for Cluster 3, containing ventral-intermediate-restricted genes. In contrast, expression in *hand2*:GFP+ cells was the strongest driver for Clusters 4 and 5, consistent with these clusters containing known and newly validated ventral-restricted genes. Consistently, the ventral-intermediate (3) and intermediate (2) clusters and the two ventral clusters (4 and 5) formed separate sub-groups when compared across all experiments. In contrast, the dorsal cluster (1) was driven by gain-of-function Notch signaling and not relative enrichment in *dlx5a*:GFP+ and *hand2*:GFP+ cells. Interestingly, expression in *edn1* mutants was a strong driver for the ventral-intermediate Cluster 3 yet disrupted intermediate Cluster 2 and ventral Cluster 4. This finding is consistent with our in situ validation showing opposite Edn1 regulation of intermediate genes *emx2* and *fsta* in Cluster 2, and regulation of *sema3bl* and *pitx1* but not *fzd9b* in Cluster 4. TOM driver analysis thus represents a powerful method for identifying new genes and regulatory mechanisms in the developing face.

### Mutational analysis reveals potential widespread genetic redundancy

We next sought to uncover potential requirements for novel domain-specific genes in zebrafish craniofacial development. To do so, we used TALEN and CRIPSR technologies to introduce early frame-shift mutations in 12 genes (*cd248a, ctgfa, ctgfb, cdh11, emx2, fsta, fstb, her6, mrrf, sfrp2, osr1, osr2*) (see Supplementary Table 2 for details) and analyzed homozygous mutant embryos for cartilage and bone defects at 5 days post-fertilization (dpf). For all mutants except *fsta* and *emx2*, no craniofacial skeletal defects were observed (data not shown). We also examined *ctgfa; ctfgb* and *osr1; osr2* double mutants yet failed to observe obvious craniofacial skeletal defects. While not displaying larval craniofacial defects, *mrrf* mutants grew slower than wild-type siblings and rarely survived past one month, and, even before general growth defects were apparent, they were unable to regenerate their tail fins (Supplementary Figure 1).

### Opposite requirements for *fsta* and *emx2* in hyoid cartilage development

Homozygous mutants for *fsta* and *emx2*, two new Edn1 targets expressed in the intermediate domain, displayed defects in the hyoid arch skeleton (Figure 6A,B). In *fsta* mutants, we detected variable alterations of the hyoid joint, a compound joint in which a small interhyal cartilage makes connections to the hyomandibular and ceratohyal cartilages on either side. The interhyal cartilage was reduced and made abnormal cartilaginous connections with adjacent cartilages, a joint fusion phenotype similar to what has been reported for *irx7*; *irx5a* mutants (Askary et al., 2015). The symplectic cartilage of *fsta* mutants was also reduced in length, and the connection between the hyomandibular and symplectic cartilages was thickened. *fsta*; *fstb* double mutants displayed a subtle enhancement of craniofacial defects compared to *fsta* single mutants (Supplementary Figure 2). In contrast, *emx2* mutants had separated symplectic and hyomandibular cartilages, with weakly Alcian-positive cells evident at the interface. As these two elements start out separate in wild types at 3 dpf (Figure 6A), we interpret the *emx2* phenotype as a failure of later cartilage fusion. These mutants also have a near complete loss of the opercular bone (a hyoid arch derivative) and abnormalities in the palatoquadrate cartilage (a mandibular arch derivative). We next tested the genetic interaction between these genes and found that expression of *fsta* is unaffected in *emx2* mutants and vice versa (Figure 6C,D). Compound *fsta*; *emx2* mutants also displayed additive phenotypes, including a shortened symplectic disconnected from the hyomandibular cartilage, loss of the opercular bone, and a variably fused hyoid joint (Figure 6A,B). Fsta and Emx2, two genes oppositely regulated by Edn1, therefore appear to act in parallel pathways to orchestrate cartilage and bone development in the intermediate domain (Figure 6E).

**Figure 6.**
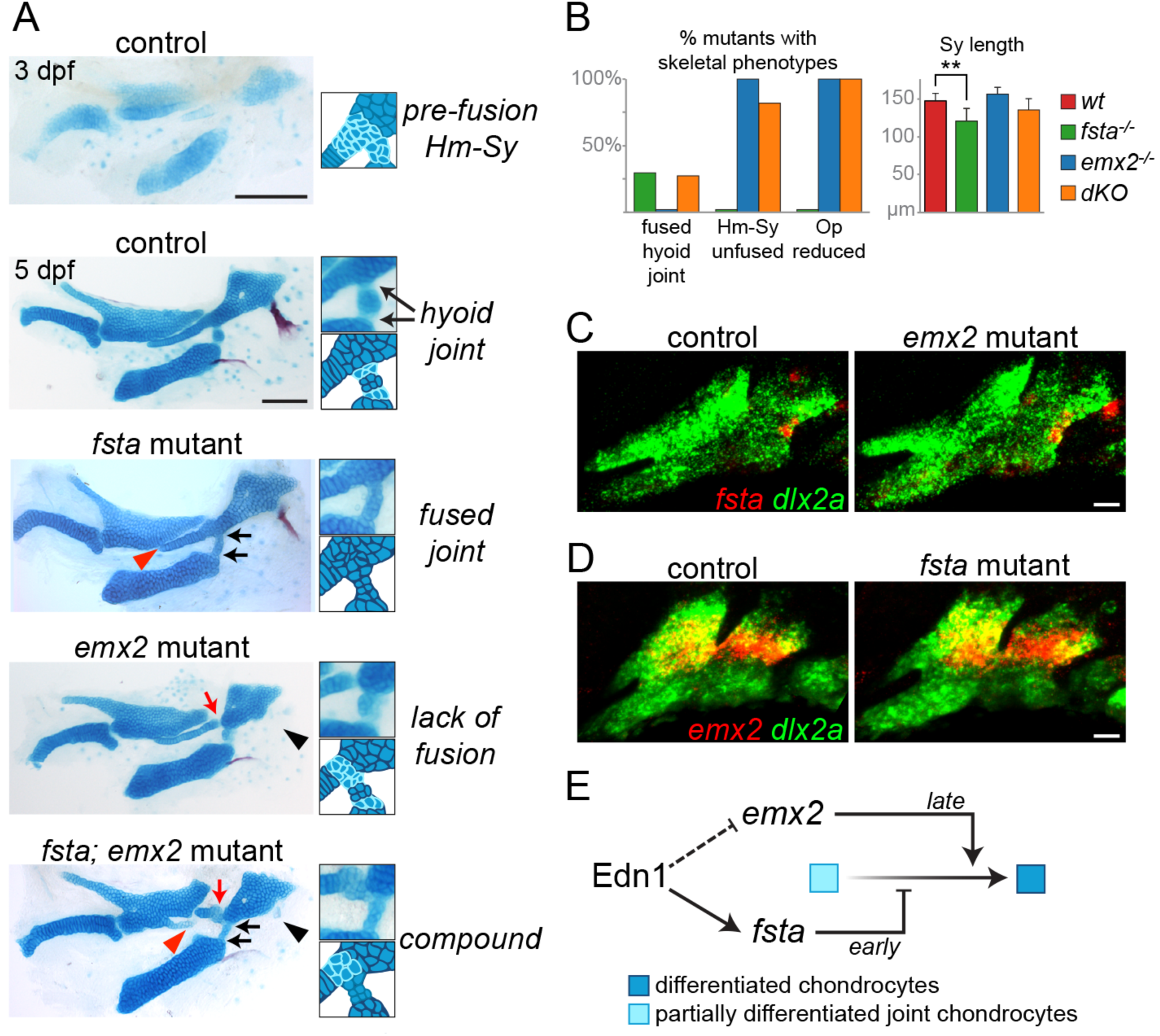
Distinct roles of Fsta and Emx2 in hyoid skeletal development. (A) Alcian Blue and Alizarin Red staining of control, *fsta, emx2* and *fsta; emx2* double mutant larvae. At 3 dpf, cells located between the hyomandibular (Hm) and symplectic (Sy) cartilages and at the forming hyoid joint region stain only weakly with Alcian Blue, reflecting their chondrogenic immaturity. By 5 dpf, rearrangements among cells at the Hm-Sy juncture have resulted in convergent extension and elongation of the Sy. The subsequent maturation of cells at the juncture fuses the Hm and Sy cartilages. Cells in the hyoid joint area remain immature and only weakly Alcian-positive. In *fsta* mutants, the Sy is shortened (red arrowhead), with a buildup of chondrocytes in the Hm-Sy juncture, and hyoid joint cells ectopically mature into differentiated chondrocytes, fusing the joint (black arrows). In *emx2* mutants, the Hm and Sy cartilages do not fuse completely (red arrow), and the opercle bone is lost (black arrowhead). *emx2; fsta* mutants (dKO) show a compound phenotype with attributes from both single mutants. Scale bar = 100 μm. (B) Penetrance of skeletal phenotypes in each genotype (57, 37 and 11 sides in *fsta^-/-^, emx2^-/-^,* and dKO) and quantification of Sy length (10, 14, 13, and 11 sides in wild type, *fsta^-/-^*, *emx2^-/-^*, and dKO). ** p < 0.0001. (C,D) Two-color in situ hybridization shows that *fsta* expression (red) is not affected in *emx2* mutants, and *emx2* expression (red) is not affected in *fsta* mutants. Arch CNCCs are co-labeled with *dlx2a* (green). Scale bar = 20 μm. (E) Model for parallel Emx2 and Fsta function in the hyoid arch. At early stages, Fsta prevents differentiation and cartilage matrix accumulation between the Hm and Sy, thus allowing chondroprogenitors to rearrange into the single stack of Sy chondrocytes. Following rearrangements, Emx2 activity may promote chondrogenic differentiation to fuse the Hm and Sy into a seamless cartilage. In contrast, there is a continuous requirement for Fsta function at the nearby hyoid joint to maintain its patency through active inhibition of chondrogenesis.

## Discussion

Our global gene expression analysis of zebrafish pharyngeal arch CNCCs has revealed general principles of arch patterning and novel expression patterns and functions of genes not previously implicated in craniofacial development. The intersection of domain-specific gene expression with changes upon signaling perturbation uncovered distinct roles for Edn1 signaling along the dorsoventral axis that may help explain the complex phenotypes of *edn1* mutants. In particular, we identified new roles for two oppositely regulated Edn1 target genes, *fsta* and *emx2,* in coordinating joint, cartilage, and bone morphogenesis in the intermediate regions of the developing arches.

### Identification of novel domain-specific arch genes

We used two complementary methods to identify co-expressed modules of genes in mandibular and hyoid arch CNCCs. The first approach took advantage of the graded expression of *hand2*:GFP and *d/x5a*:GFP transgenes along the dorsoventral axis to group genes into four compartments, and the second approach mined co-variation across 19 RNAseq data sets to identify genes with similar expression patterns and/or regulation. A limitation of the first strategy is that filtering thresholds are empirically determined, and that genes must pass all thresholds to be included (which likely accounts for some known genes, such as *dlx6a,* being excluded). Empirical shifting of thresholds, guided in part by anchoring well-characterized genes in each cluster, led to a balance between the number of false positives (genes not expressed in the predicted domains) and false negatives (genes with known domain-specific expression not included on the list). An advantage of the second co-variance strategy is that it is unbiased, although both the relative enrichment in *hand2*:GFP+ and *dlx5a*:GFP+ domains and expression changes in response to signaling perturbation drive clustering. Nonetheless, considerable concordance between the approaches points to the validity of each. For example, four ventral genes (*pitx1*, *fzd9b*, *foxf1*, *foxj2a*), four ventral-intermediate genes (*dlx3b, dlx4b, dlx5a, msxe*), and five intermediate genes (*emx2, fsta, igfbp5b, grem2b, foxd1*) were similarly identified by *hand2*:GFP*/dlx5a:*GFP filtering and co-variance analysis. Using these types of analyses, we uncovered a number of new genes with validated domain-specific expression. In the ventral domain, these included the S-phase kinase-associated protein 2 (*skp2*) and the mitochondrial ribosome recycling factor (*mrrf*), perhaps reflecting distal growth of this domain to elongate the lower jaw (Bonilla-Claudio et al., 2012; Medeiros and Crump, 2012). In the ventral-intermediate domain, we uncovered specific expression of Fgf binding proteins (*fgfbp2a* and *fgfbp2b*), suggesting fine regulation of Fgf signaling in this domain. In the intermediate domain, we discovered two putative Bmp inhibitors with tightly restricted expression near the developing hyoid joint, consistent with prior data showing complex regulation of Bmp signaling being important for joint specification (Salazar et al., 2016; Smeeton et al., 2017). In the dorsal domain, we uncovered selective expression of genes previously implicated in earlier neural crest and ectomesenchyme development, including Snail transcription factor (*snai1a*) (LaBonne and Bronner-Fraser, 2000), Cadherin 11 (*cdh11*) (McLennan et al., 2015), potassium channel tetramerization domain-containing 15a (*kctd15a*), which interacts with *tfap2a* (Zarelli and Dawid, 2013), and endosialin (*cd248a*) (Das and Crump, 2012); this signature is consistent with our previous findings that dorsal arch CNCCs differentiate later than other arch CNCCs (Barske et al., 2016). We also, however, found some false positive genes, in particular in the category annotated as dorsal. The secreted frizzled-related protein 2 (*sfrp2*), whose mouse homolog was reported to be upregulated in *Dlx5/6* mutants, consistent with dorsal enrichment (Jeong et al., 2008), was restricted to the dorsal arches in zebrafish yet confined to the mesoderm. Similarly, expression of Ras-associated domain family member 10a (*rassf10a*) was largely in arch ectoderm and not CNCCs. We cannot rule out lower levels of expression in CNCCs, or some rare non-CNCC arch cells being included in our FACS-sorted populations. Future generation of transgenes to specifically label dorsal CNCCs, as well as other domains such as the frontonasal and maxillary prominences, should aid in creating more precise expression atlases of additional arch regions.

### Region-specific roles of Ednl and Jagged-Notch signaling in arch patterning

Previous work had suggested greater roles for Edn1 signaling in intermediate versus more ventral domains (Alexander et al., 2011; Zuniga et al., 2011), and for Jagged-Notch signaling in the dorsal arches (Zuniga et al., 2010). By analyzing how gene modules of distinct arch domains are affected by signaling perturbations, we confirm this on a genomic scale (summarized in Fig. 4E). A more prominent role for Edn1 in controlling gene expression in intermediate mandibular and hyoid arch domains, which generate joints and the palatoquadrate and symplectic cartilages, helps explain why these skeletal elements are most sensitive to partial reduction of Edn1 function (Miller and Kimmel, 2001) and mutation of its downstream effectors Plcb3 and Mef2ca (Walker et al., 2006; Walker et al., 2007). Conversely, the ventral-most elements of the mandibular and hyoid arches, such as the basihyal, are spared in severe *edn1* mutants (Miller et al., 2000), consistent with the expression of most ventral genes in this study (*smad6a, skp2, fzd9b*) and previous reports (*satb2*) (Sheehan-Rooney et al., 2013a) being unaffected by Edn1 perturbations. However, some ventral genes, such as *hand2* and *pitx1*, are lost in *edn1* mutants, although regulation of *hand2* may be indirect through the Edn1 targets Dlx5/6 (Miller et al., 2003; Yanagisawa et al., 2003). Edn1-independent ventral genes may instead depend on Bmp signaling. Indeed, *Smad6* and *Satb2* were identified as direct targets of Bmp-dependent pSMADs in mice (Bonilla-Claudio et al., 2012), and *satb2* is a target of Bmp signaling in zebrafish (Sheehan-Rooney et al., 2013a). In the dorsal domain, we also found only a subset of genes to be regulated by Jagged-Notch signaling (e.g. *cd248a, pou3f3a,* and *pou3f3b* but not *cdh11* and *kctd15a*), consistent with the relatively mild dorsal phenotypes of *jag1b* and *notch2*; *notch3* mutants (Zuniga et al., 2010; Barske et al., 2016) and suggesting Notch-independent regulation of some aspects of dorsal identity. Whereas we found generally good correspondence between changes in RNAseq values and in situ validation in *edn1* and *jag1b* mutants, in situ validation but not RNAseq revealed differences in *stmn1a, sema3bl,* and *cdh11* expression in *edn1* mutants. As *stmn1a* and *sema3bl*show broad arch expression, profiling all arch CNCCs likely dilutes the effect of selective loss of their ventral expression domains in mutants. Likewise, a shift of *cdh11* expression from dorsal to ventral domains in mutants would not necessarily result in a total expression difference throughout arch CNCCs. These findings suggest that examining expression changes in CNCCs sorted from distinct arch domains in animals with signaling perturbations may be a better way to detect how signaling affects expression patterns.

### A lack of obvious craniofacial phenotypes in mutants for many arch-specific genes

The ease of genetic manipulation makes zebrafish an attractive system for performing reverse genetic analysis of craniofacial development. However, homozygous loss-of-function mutants for only two of the 12 domain-specific genes tested (*emx2* and *fsta*) showed clear facial cartilage and/or bone phenotypes in larvae. There are several possible explanations for the lack of observable phenotypes. First, although we selected mutations causing premature translational termination before critical conserved domains, it is possible that some mutations do not create true nulls. Second, some mutants may have craniofacial defects that we failed to appreciate, for example in other arch derivatives such as ligaments or long-lived progenitors. Third, maternal contribution of mRNA and/or protein could compensate for zygotic loss-of-function. In some cases (e.g. *ctgfa*), maternal-zygotic null mutants did not display larval craniofacial defects. For *mrrf,* the growth delay and tail fin regeneration defects of *mrrf* mutants could be explained by depletion of remaining maternal stores, similar to what has been reported for other mutants in mitochondrial proteins (Rahn et al., 2015). It thus remains possible that *mrrf* expression in the ventral-most arches reflects rapid growth and/or metabolism of this domain. Fourth, there may be genetic compensation (Rossi et al., 2015), although *ctgfa*; *ctgfb* and *osr1*; *osr2* compound mutants had no apparent larval craniofacial defects. Large-scale mutational screens in zebrafish have found a surprisingly small number of genes required for larval viability (~6%), suggesting a high degree of genetic redundancy in zebrafish (Kettleborough et al., 2013). In addition, the identification of multiple alleles for craniofacial mutants suggests that previous screens are approaching saturation for obvious larval skeletal defects (Neuhauss et al., 1996; Piotrowski et al., 1996; Schilling et al., 1996; Nissen et al., 2006). Our findings may therefore indicate that many of the single gene mutants with obvious craniofacial patterning defects in zebrafish may already have been found.

### Complex regulation by Edn1 coordinates intermediate arch morphogenesis

A curious feature of *edn1* mutants, as well mutants for its effector *mef2ca*, is the phenotypic variability of intermediate domain-derived skeletal elements, including joints and the opercle bone (Kimmel et al., 2003; DeLaurier et al., 2014). Our analysis of two newly identified Edn1 target genes, *emx2* and *fsta,* may shed some light on this variability. For example, the gain or loss of the opercle in *edn1* mutants might reflect the observed variability in *emx2* upregulation, which we find to be required for opercle formation. The loss of the hyoid joint and reduction in symplectic cartilage in *fsta* mutants are also similar to what is seen in Edn1 pathway mutants (Miller et al., 2000; Walker et al., 2006; Walker et al., 2007; DeLaurier et al., 2014), consistent with our finding that intermediate domain *fsta* expression is lost in *edn1* mutants. Our previous analysis of similar phenotypes in *irx7*; *irx5a* compound mutants showed that inappropriate chondrogenic differentiation at the juncture between the nascent symplectic and hyomandibular cartilages prevents these cells from rearranging and thus lengthening the symplectic (Askary et al., 2015). In contrast, the hyomandibular and symplectic cartilages fail to connect in *emx2* mutants. In one model, temporal regulation by Edn1 in the intermediate domain may result in Fsta blocking cartilage differentiation at early stages to allow symplectic formation, and then Emx2 promoting cartilage differentiation at later stages to fuse the symplectic and hyomandibula into a seamless cartilage (Figure 6E). Interestingly, *Emx2* mutant mice lack the incus cartilage of the middle ear (Rhodes et al., 2003), which is homologous to the palatoquadrate affected in fish *emx2* mutants. Part of the arch patterning function of Emx2 may thus be conserved from fish to mammals. Our transcriptome-driven analysis of arch regionalization has therefore provided new insights into how Edn1 signaling regulates a delicate balance of cartilage differentiation to fine-tune skeletal shape.

## Materials and Methods

### Zebrafish lines

The University of Southern California Institutional Animal Care and Use Committee approved all experiments on zebrafish (*Danio rerio*). Published lines include *Tg*(*hand2:eGFP*) (Kikuchi et al., 2011), *dlx5a^j1073Et^* (hereafter *dlx5a:GFP,* (Talbot et al., 2010)), *Tg*(*fli1a:eGFP)^y1^* (Lawson and Weinstein, 2002), *Tg(sox10:DsRed-Express)^el10^* (Das and Crump, 2012), *Tg(sox10:GFPCAAX)* (Askary et al., 2015), *sucker/edn1^tf216^* (Miller et al., 2000), and *jag1b^b1105^* (Zuniga et al., 2010). Mutant lines were genotyped as previously described.

### Fluorescence activated cell sorting and RNA sequencing

*fli1a:GFP*; *sox10:DsRed* and *hand2:GFP*; *sox10:DsRed* fish were incrossed to generate embryos, and *dlx5a:GFP; sox10:DsRed* was outcrossed to avoid homozygosity of the *dlx5a^j1073Et^* insertional allele. Embryos were dissociated as previously described (Covassin et al., 2006), with minor modifications (Barske et al., 2016). Cells were sorted based on GFP and DsRed expression on a MoFlo Astrios instrument (Beckman-Coulter, Brea, CA, USA) into RLT lysis buffer (Qiagen, Hilden, Germany), and total RNA was extracted using the RNeasy Micro kit (Qiagen). RNA integrity was assessed on Bioanalyzer Pico RNA chips (Agilent, Santa Clara, CA), cDNA synthesized with the SMART-Seq Ultra Low Input RNA Kit (Clontech, Mountain View, CA), and libraries generated with the Kapa Hyper prep kit (Kapa Biosystems, Wilmington, MA) and NextFlex adapters (Bioo Scientific, Austin, TX). 75 bp paired-end sequencing was performed on a NextSeq 500 machine (Illumina, San Diego, CA).

### RNAseq data analysis and statistical tests

After trimming using Partek Flow default criteria, sequencing reads were aligned to zebrafish GRCz10 (Ensembl_v80) using TopHat 2 algorithm. Aligned reads were quantified using the Partek E/M algorithm and normalized to yield TPM values, controlling for sequencing depth disparities across samples (Wagner et al., 2012). Data are accessible through GEO Series accession number 18301072. To test whether log_2_FC values for each group of genes were significantly different from zero (Figure 3), we used the Shapiro-Wilk test for normality to determine whether a one-sample t-test or a Wilcoxon signed-rank test was appropriate. The Bonferroni correction was then applied on the resulting *p* values of one-tailed tests to account for multiple comparisons. The Mann-Whitney U test for two independent samples was performed in MS Excel 2016 using the Real Statistics Resource Pack software (release 4.9; www.real-statistics.com) to compare effects of Edn1 and Notch signaling, as the data from at least one group were not distributed normally.

### Co-variance analysis

A weighted gene co-expression network analysis (WGCNA) (Langfelder and Horvath, 2008) was run on *m* genes exhibiting the highest variance across *n* samples (*m =* 6000, *n =* 19), yielding an *m* * *m* Topological Overlap Matrix (TOM) that links genes by correspondence of correlated genes. The exponent *β* was selected to yield scale-free topology as defined by minimum power required to output maximal R^2^. The TOM Driver Array (TDA) was computed by taking the average TOM value across genes of interest, then deriving the deviation in TOM when each sample was removed.

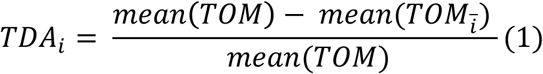

These values are normalized to produce nTDA, which spans [-1,1].

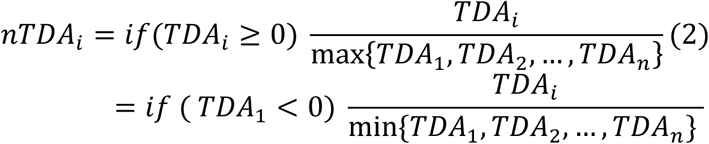

Samples with a positive TDA drive clustering, while samples with negative TDA disrupt clustering.

### In situ hybridization and immunohistochemistry

Partial cDNAs were PCR-amplified with Herculase II Fusion Polymerase (Agilent Technologies, Santa Clara, CA), cloned into pCR_Blunt_II_Topo (ThermoFisher Scientific, Waltham, MA), linearized, and synthesized with Sp6 or T7 RNA polymerase (Roche Life Sciences, Indianapolis, IN) as specified (Supplementary Table 3). In situ hybridization was performed as published (Zuniga et al., 2010), costaining with *dlx2a* (Akimenko et al., 1994) or rabbit anti-GFP antibody (Torrey Pines Biolabs, Secaucus, NJ) to highlight arch CNCCs. Imaging was performed with a Zeiss LSM800 confocal microscope and presented as optical sections or maximum intensity projections as specified. Approximately 6-10 controls and 3-7 mutants were imaged for each probe.

### Mutant generation and skeletal staining

Twelve mutant lines were created via TALEN (Sanjana et al., 2012) or CRISPR/Cas9 (Jao et al., 2013) mutagenesis as described (Barske et al., 2016). Germline founders were detected by screening their F1 progeny by restriction digestion of PCR products, followed by sequencing to identify frameshift indels (Supplementary Table 2). Alcian Blue and Alizarin Red staining of cartilage and bone was performed as described (Walker and Kimmel, 2007). Symplectic cartilage length was measured with ImageJ and compared with unpaired t-tests.

## Acknowledgments

We thank Megan Matsutani and Jennifer DeKoeyer Crump for fish care, Lora Barsky and Jeffrey Boyd at the USC Stem Cell Flow Cytometry Core Facility for FACS, Charles Nicolet at the USC Norris Cancer Center Molecular Genomics Core for sequencing, and Yibu Chen for help with sequencing analysis.

